# ELICE16INDURES ^®^: a plant immune-priming activator targeting jasmonate metabolism via TIFY and RHOMBOID proteins in *Hordeum vulgare*

**DOI:** 10.1101/2021.08.11.455979

**Authors:** Géza Hegedűs, Márta Kiniczky, Ágnes Nagy, Péter Pekker, Balázs Lang, Lajos Gracza, József Péter Pallos, Zsófia Thomas-Nyári, Kincső Decsi, Barbara Kutasy, Kinga Székvári, Ákos Juhász, Eszter Virág

## Abstract

Priming activity of plant-based allelochemicals is advanced research nowadays meaning a high potential in sustainable agriculture. The ELICE16INDURES^®^ (RIMPH LTD, Hungary) plant conditioner of CO_2_ botanical extracts is rich in plant-active ingredients such as phenolic compounds, alkaloids, and flavonoids formulated in small multilamellar liposomes. This product was investigated *in autumn barley (Hordeum vulgare)*. Field experiments of ELICE16INDURES showed augmented NDVI values interconnected with higher photosynthetic activity and yield increase. Background of the better vitality of plants was investigated by whole genomic gene expression profiling and showed an enhanced response to wounding, jasmonic acid, oxidative detoxification, and chloroplast activity. Among top50 differentially expressed genes the TIFY domain protein TIFY11B and RHOMBOID-like protein 2 related to JA signaling were up-regulated in field-collected samples. Phytotron experiments of barley were set up to validate and evaluate the transcriptomic effect of ELICE16INDURES. Well-studied priming active agents such as salicylic acid and beta-aminobutyric acid were compared with ELICE16INDURES and confirmed as priming inducer material with positive regulation of TIFY11B, TIFY3B, TIFY9, TIF10A, and RHOMBOID like protein 2 by using NGS GEx and RT-qPCR methods.

**One-sentence summary:** ELICE16INDURES^®^ is a plant conditioner agent with a high amount of allelochemicals encapsulated into small multilamellar liposomes and found as an immune priming activator tested in *H. vulgare* field and phytotron cultures.

## INTRODUCTION

The application of natural-based activators is the most innovative and ecologically safe resolution to increase crop vitality and production. Nowadays sustainable agricultural production and preservation of biodiversity in agricultural fields pay great attention worldwide therefore the development of alternative plant protection is in focus avoiding the use of chemicals: pesticides, nutrients, or soil improvers (Du Jardin, 2015). Many companies are developing products of biologically active agents (Calvo et al., 2014; Sharma et al., 2014) based on plant signaling molecules so-called "metabolic enhancers” or “biostimulants” among which natural vitamins, phytohormones, amino acids, and their derivatives are the most significant (Ugena et al., 2018). To improve yield and abiotic stress tolerance in agriculturally important cereal cultures the use of biostimulants is a promising approach, however, less applied in practice than synthetic ones.

The responses to biostimulants were first studied in *Arabidopsis thaliana* (*A. thaliana*, mouse-ear cress) because this plant has one of the smallest plant genomes (Toscano et al., 2018; Geelen and Xu, 2020). In this experiment, 127 genes showed at least 3-fold activity against an untreated plant under the influence of a commercially available biostimulant formulation. Selivanova et al. (2015) investigated the physiological processes of cucumber plants under the influence of several commercially available biostimulant products. The results of the experiment showed an increased plant metabolism as a result of the treatment, which in turn increased yields. Klokić et al. (2020) examined the effect of biostimulants on tomato yield characteristics. As a result of the experiment, it can be concluded that the treatments with 10 different biostimulants influenced the characteristics of the fruits, such as the total phenol and flavonoid content and the total antioxidant capacity (Klokić et al., 2020). Špoljarevic et al. examined the activity of antioxidants in strawberry (*Fragaria x ananassa Duch*) leaves in two biostimulant treatments with two nutrient replenishment. The specific activities of guaiacol peroxidase (GPX), catalase (CAT), ascorbate peroxidase (APX) and glutathione reductase (GR) were investigated in leaves. Based on the results, it may be concluded that the values of antioxidant enzymes were higher in the leaves after treatment with biostimulants and reduced nutrient replenishment. APX and GR showed the highest activity in the treated plants, and the strongest relationship was also between these two enzymes, which are enzymes of the ascorbate-glutathione cycle (Špoljarević et al., 2010). Later they summarized the researches on different biostimulants in horticultural plant species with the conclusion of the effects of physiologically active compounds on plants may depend on dose, time of treatment, growth conditions, and plant species. Therefore they suggest intensive research in this field (Parađiković et al., 2019).

Some biostimulants may protect plants from a broad range of pathogens by activating the plant immune system (Burketova et al., 2015; Vargas-Hernandez et al., 2017). The effect of these materials lies in not targeting pathogens like pesticides or not directly induce immune response overcoming by pathogenic microbes however, potentiate plant defense mechanism long term, therefore they are called priming-active or imprimatin-compounds (Noutoshi et al., 2012). From the point of view of field usage of biostimulants, priming-active materials are the more cost-effective since the induced resistance may be triggered and optimized. Some natural and synthetic compounds have demonstrated good priming-inducing activity in laboratory and field conditions, such as the nonprotein β-aminobutyric acid (BABA) or the phytohormone salicylic acid (SA) (Beckers and Conrath, 2007; Walters et al., 2014). The effect of priming-active elicitors on protection and signaling pathways has already been demonstrated, so its potential use in the field is well-founded and highlighted. From the 20 years of the discovery of these compounds, the mechanisms of plant immune responses are also gaining more and more light and paving the way for new plant protection strategies (Schwessinger and Ronald, 2012).

Several plant extracts were used as priming agents among which the seed priming gain in more attention in field crops (Farooq et al., 2019). In these experiments the allelochemicals such as phenolic compounds, alkaloids, terpenoids, flavonoids, saponins, and steroids may act beneficially to plant growth, germination reaching higher vitality and resilience (Narwal, 2006; Adeniyi et al., 2010; Narwal, 2012; Moses et al., 2014; Mujeeb et al., 2014; Latif et al., 2020). For example, the reduction in mortality and high seedling vigor in tomatoes was reported by primed tomato seeds with Azadirachta, Chlorophytum, and Vinca (Prabha et al., 2016). Saponins can enhance nutrient absorption as they are readily soluble in water. Alkaloids, saponins, and phenolic compounds present in the leaves of various plants are involved in the production of antioxidant activities and protect the plants against pathogens (Satish et al., 2007). The strong presence of the above-mentioned phytochemicals was proven in different extracts of garlic cloves (Divya et al., 2017) and their antimicrobial properties were also investigated against pathogens such as *Fusarium* and *Aspergillus* species (Olusanmi and Amadi, 2010).

An advanced method in priming is the use of nanoparticles in the range of 100-500 nm in size. Although this technology is in little attention, there are some reports on the positive impact of their uptake and nutrient delivery applied in foliar and seed priming as well (Dutta, 2018; Abdel-Aziz et al., 2019). However, the priming with nano-size liposomes as macromolecule carriers was not yet been reported in plants. Some studies investigated the use of liposomes to supplement plant growth and overcome acute nutrient deficiency (Karny et al., 2018). The advantage of liposome encapsulation is to reduce the quantity of bioactive compounds to reach the same impact. The product ELICE16INDURES^®^ plant conditioner (Research Institute for Medicinal Plants and Herbs Ltd. Budakalász, Hungary) contains Garlic cloves CO_2_ extracts, and a high amount of above-mentioned allelochemicals encapsulated in 250-350 nm size multilamellar liposomes.

In this study, the transcriptomic effect of ELICE16INDURES was investigated and compared with BABA and SA priming agents in the field and phytotron experiments on autumn barley *(Hordum vulgare)*. Chloroplast activity predictions were performed by remote sensing in field cultures. Monitoring the extension of photosynthetic surface in barley cultures by using agro-drone a higher vitality was detected in the ELICE16INDURES treated plots that were manifested also in augmented crop yield.

## RESULTS AND DISCUSSION

### Testing of ELICE16INDURES in field experiments

#### Determination of NDVI in field populations of *H. vulgare*

The NDVI value provides information on the photosynthetic activity of plants (Kalbi et al., 2014), i.e. it examines how much of the light in the photosynthetic wavelength range can be used by the plant. NDVI gives a picture of the differences in vegetation activity and vegetation rate within the field. This provides useful information for assessing heterogeneity within the field and may draw attention to the occurrence of any problems or changes within the field (inland water, prolonged germination, nutrient deficiency, wildlife damage and plant disease). The development of vegetation can be followed along with the development of NDVI in each field, the effect of different varieties, hybrids, or different crop production technologies can be compared. True, these are still not exact values, but they can provide support. In this case, NDVI provides information on the effect of ELICE16INDURES. Using NDVI spectral analysis we monitored the area as indicated in Figure 2A.

**Figure 1.**
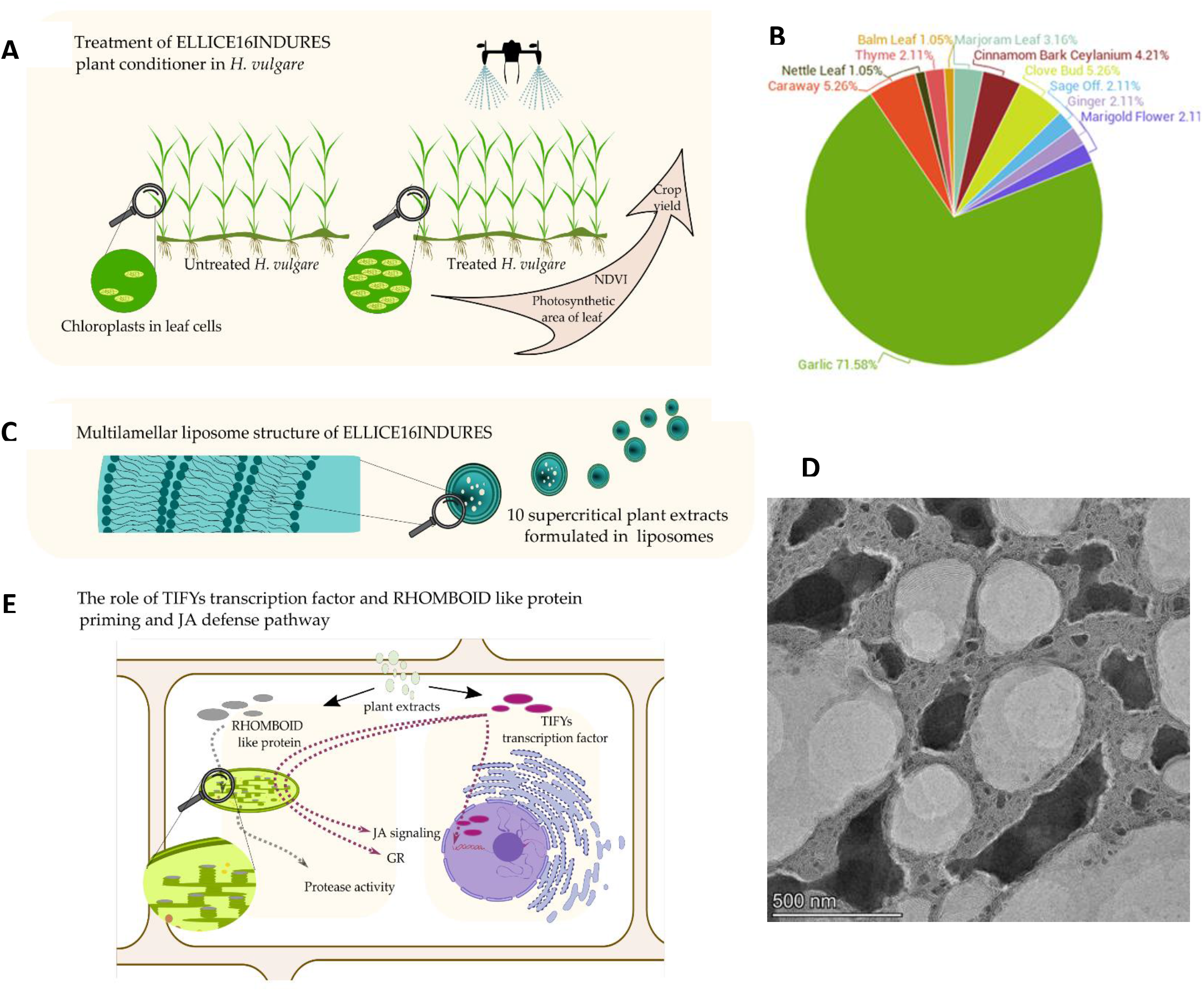
Summary of ELICE16INDURES^®^ structure and mode of action. Field treatment of ELICE16INDURES plant conditioner with drone spraying **(A)**. Higher chloroplast and photosynthetic activities were monitored by NDVI calculation. The main component of ELICE16INDURES is the garlic CO_2_ extract which ratio is more than 70% in the liposomes **(B)**. Multilamellar small liposome structures **(C, D)** helps the delivery of the priming active material complex to the plant cell. According to transcriptomic profiling, TIFY domain proteins and RHOMBOID-like proteins are induced and lead to augmented protease, glutathione reductase activity, and Jasmonate signaling **(E)**.

**Figure 2.**
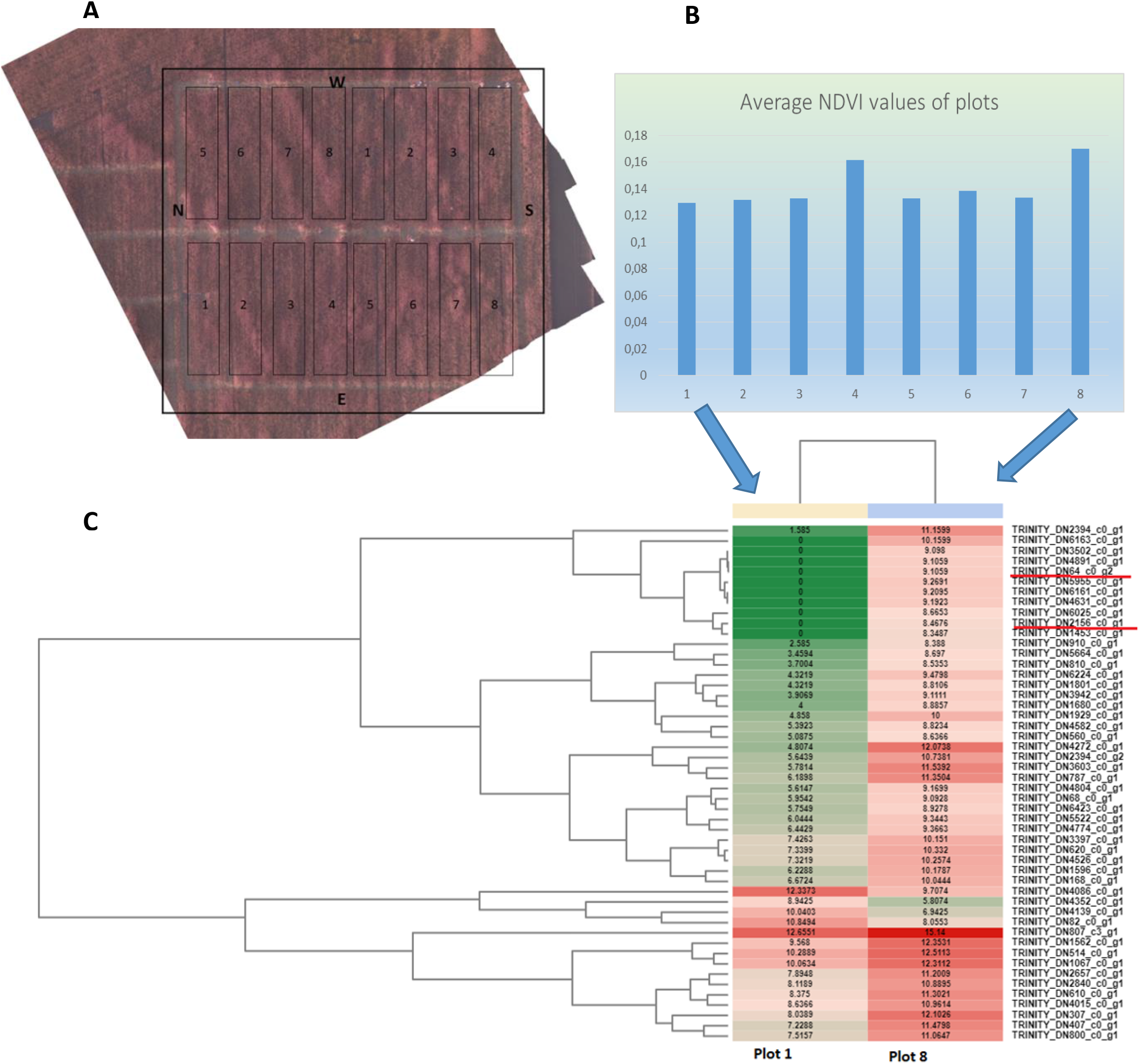
Field experiments of ELICE16INDURES^®^ on autumn barley cultures. Plots and doses of ELICE16INDURES^®^ applied in field experiments: **1**, control without treatment; **2**, positive control treated with Fitokondi^®^; **3**, 10 g/ha; **4**, 20 g/ha; **5**, 30 g/ha; **6**, 60 g/ha; **7**, 120 g/ha ; **8**, 240 g/ha ELICE16INDURES^®^. Experimental areas are visualized by a combined photograph from drone photos **(A)**. A near-infrared camera was used to calculate NDVI from each pixel. The average NDVI of plots calculated from pixel values is visualized on figure **(B)**. Heatmap of top 50 DEGs of *H. vulgare* treated by ELICE16INDURES^®^ in the field **(C)**. Samples were collected randomly from the plot **8** and **1**. Genes highlighted with red are involved in JA-pathway: TRINITY_DN64_c0_g2, protein TIFY 11e-like and TRINITY_DN2156_c0_g1, RHOMBOID-like protein 2. Annotation of heat map see in Supplemental Table S2.

Photosynthetic traits are indicators of the physiological status of plants. The higher NDVI value suggested a larger area with enhanced photosynthetic activity in leaves of investigated barley culture. This state might have resulted in fewer patients or better assimilation of nutrients, water, or light. We hypothesized that the higher NDVI values correlated with the healthier leaf surface. Therefore we chose plot 8 and plot 1 to highlight the background of this phenomenon. NGS sequencing (GEx) was carried out using plant materials collected in the given plots. We performed bioinformatics analysis and determined the top50 DEGs (Figure 2C).

#### Determination of DEGs in field populations of *H. vulgare*

NGS GEx was performed from randomly collected leaves grown in the two parallel plots (see Figure 2). Three biological repeats were investigated in both conditions. Among the top50 DEGs the most interesting genes are detailed in Supplemental Table S1. Results highlighted an outstanding defense response interconnected with JA pathway and oxidative regulation. TIFY 11 –like proteins have a role in response to wounding and JA-mediated immune response. TIFY11 is a member of the family TIFY with a conserved domain taking part in jasmonate signaling. These proteins may be up-regulated by jasmonate, wounding, and herbivores. Chung and Howe report on the critical role of the TIFY motif in repression to JA signaling by a stabilized splice variant of the JASMONATE ZIM-domain protein JAZ10 in *A. thailana* (Chung and Howe, 2009). Our results indicated a strong up-regulation of TIFY11 gene after two days of ELICE16INDURES treatment in field conditions. This may suggest that repression of JA pathway was eliminated by TIFY11 protein under ELICE16INDURES conditions, presumably. This result points out the priming-activity of ELICE16INDURES which may be associated with JA pathway regulation. JAZ protein family shares a conserved TIFY motif within the ZIM domain which are key regulators of jasmonate signaling (Thines et al., 2007). Interestingly we found a strong up-regulation of RHOMBOID-like protein 2 interconnected with TIFY 11. Rhomboid proteases regulate intramembrane proteolysis recognizing a distortion in a short stretch of amino acids in a transmembrane region (Kanaoka et al., 2005). This function is a fundamental mechanism for controlling a wide range of cellular functions. The relationship of rhomboid proteases with JA pathway was described by (Knopf et al., 2012). They report on RHOMBOID function, with chloroplast rhomboids being involved in allene oxide synthase (AOS) accumulation in the *A. thailana* inner envelope membrane which is one of the three key enzymes of the chloroplastic step of JA biosynthesis (Figueiredo et al., 2015). We found that rhomboids were up-regulated in this experiment suggesting that JA metabolism was strengthened at the peroxisome-related step involving up-regulation of peroxidases as well.

#### Testing of ELICE16INDURES in phytotron experiments

Exogenous treatment of SA, BABA, ELICE16INDURES, and their combinations were performed according to the experimental design (Figure 3 A B). Impact on the whole genome expression profiling of ELICE16INDURES was in focus comparing with SA and BABA well studied immune-priming activators. The combination of the investigated three agents resulted in 16 samples, that gene expression comparison was helped by mathematical modeling of most different samples and genes.

**Figure 3.**
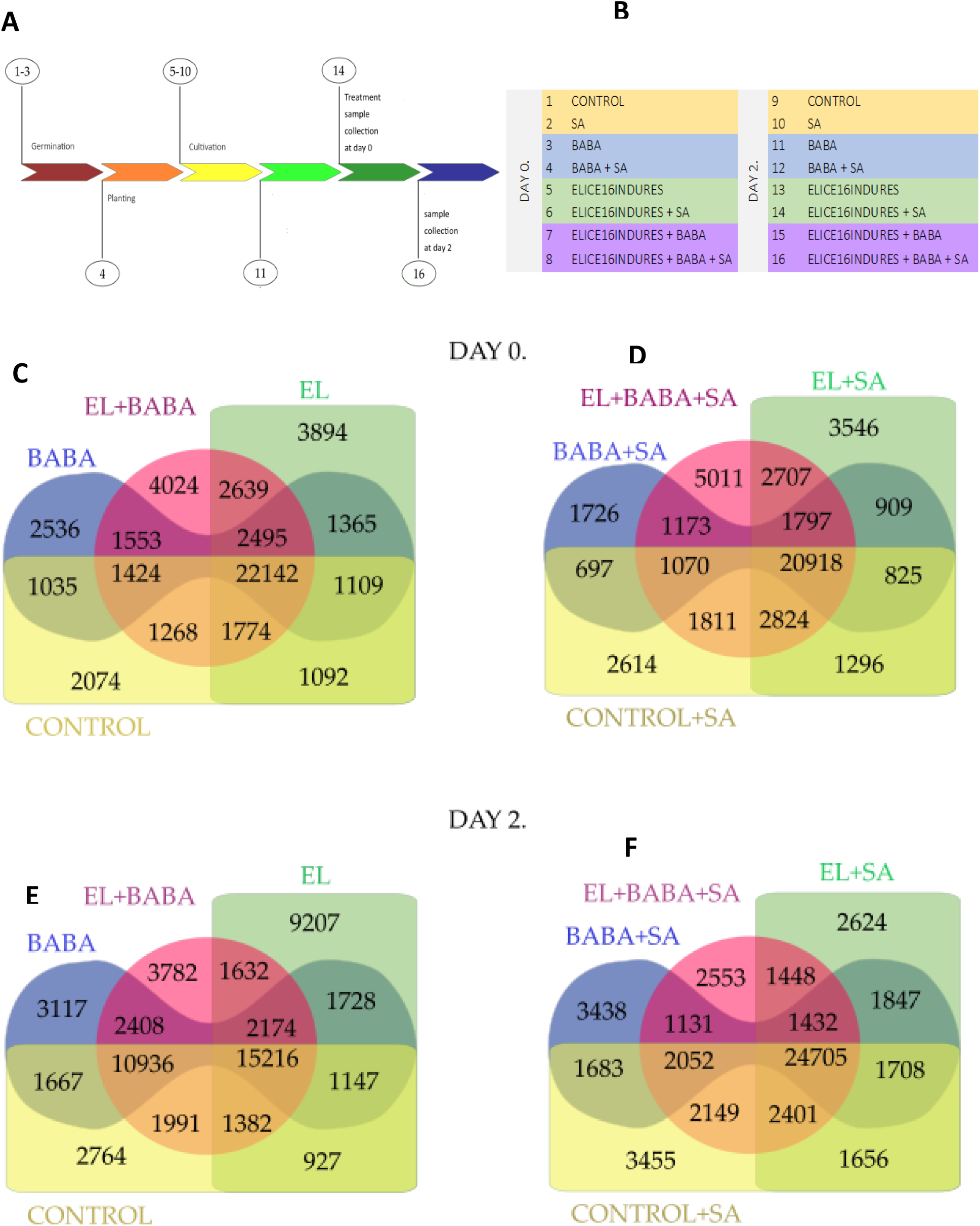
Venn diagrams. Numerical analysis of individually and commonly expressed genes in *H. vulgare* treated by priming-active materials. Experimental design of phytotron experiments **A** and **B**. Abbreviations: EL, ELICE16INDURES; SA, salicylic acid; BABA, beta-amino butyric acid. Treatments of priming inducing agents were performed on day 14. Sample collections were performed at day 14 **(Day 0)** at day 16 **(Day 2)**. Samples were collected at day 0 **(C, D)** and 2 **(E, F)** after treatments. Control samples were not treated, these were cultivated under the same conditions as the treated samples. For all 16 samples, 4 × 4 groups were determined to visualize the numerical differences between the expressed genes. **C** and **E** diagrams compare numerically the genes expressed in the case of El and BABA treatments and their combinations. **D** and **F** tween diagrams compare numerically the genes expressed between the treatments EL and BABA supplemented with SA.

#### De novo assembly and mapping

The reference transcript dataset (TrinityH16-transcript, Supplemental Data Set S1) contained 73,301 nucleotide sequences (contigs) that average length was 286 bases with minimum and maximum length 83 and 1,918 bases.

All reads from each biological repeat (3 × 16) were separately mapped against the reference dataset one and multiple times. Statistic of mapped reads was performed and compared between all replicates. No significant differences in transcript level and mapping were found between the biological replicates. Therefore reads derived by the same conditions were captured as one sample. The combined read sets were then aligned to the reference containing transcripts. Using this dataset we collapsed unique and common sequence regions among isoforms into a single linear sequence. In our system, this step allows us to analyze the *“in silico poled”* replicates as one sample. If there is a difference in the coverage complexity of a given transcript due to the replication, can be used as average information from the given transcript, representing a gene. The SuperTranscript (assembly at gene level) (TrinityH16-supertranscript, Supplemental Data Set S2) contained 60,614 nucleotide sequences, which were further analyzed.

#### Functional annotation

Functional annotation of TrinityH16-supertranscript reference dataset was performed. Annotation results are summarized in Supplemental Table S2 that resulted in 58.65 % blast results in the NCBI nr database.

#### GO categories of the reference transcriptome

The entire annotated data set (TrinityH16-supertranscript) indicated an approximate functional distribution of GO term categories. To get the most comprehensive information, 2 and 7 levels of categories were taking into account (Supplemental Figure S1). The number and category distribution indicated a high complexity degree of the data set, which allows further GO investigations. Annotation results were used to perform expression comparison during sample-by-sample and gene-by-gene analysis. The high complexity of GO categories correlated with the complexity of reference transcriptome which base the further comparisons of samples involved in this dataset.

#### Determination of gene expression profiling in 16 samples

The following relations were determined based on count table RPM values: (i) the number of individually expressed genes; (ii) samples having the largest differences in the overall transcription profile; (iii) genes having the greatest changes; (iv) differential expressions of selected samples.

#### Individually expressed transcripts

Transcripts only expressed in the given sample -were determined based on the count table data. For all 16 samples, 4 × 4 groups were determined to visualize the numerical differences between the expressed genes. The number of common and unique elements of sets are depicted on Venn diagrams (Figure 3). For better understanding grouping was based on the treatment complexity in the 0 and 2 days.

From the number of induced genes, results showed that on day 2, the level of gene expression was more significant in the treated samples, except for the ELICE16INDURES + BABA combination treatment, where the number of individually expressed genes decreased by the second day. Based on the control sample, on day 2, the SA stimulus-induced alterations in the expression of more than 1000 genes. This trend was also observed by BABA treatments. Combined SA + BABA stimulus may cause a greater change in gene expression than alone. However, this phenomenon showed the opposite in the case of ELICE16INDURES. These samples showed a larger change without SA stimulus. Here, the number of unique transcripts stands out dramatically on day 2. Dual treatment caused fewer transcriptomic changes in each case than the treatments alone.

#### Searching of sample differences by 2-norm

We used this method to determine which treatment might cause the largest change in the number of differently expressed genes. Treatment (sample) combinations giving the greatest distances according to the 2-norm were examined. We focused on the outlier values, therefore larger values are marked in green and smaller values in red (Figure 4C). Pink indicates the top 10 combinations (which showed the most larges changes). Based on the distance values in the matrix we selected 2 × 3 samples, which was analyzed in pairwise analysis: (i) 1 vs 10 (ii) 1 vs 11; (iii) 1 vs 13; (iv) 1 vs 15; (v) 1 vs 16; and the outstanding (vi) 9 vs 8 combinations. Taking into account the comparison of different effects of treatments alone and in combinations we grouped the pairs into two categories. The direct effect of ELICE16INDURES, BABA, and SA (i-iii) and combined effects of them (iv-vi). Pairwise differential expression analysis with the selected samples was performed. Based on this data the following conclusion might be performed: SA treatment caused the largest change among the investigated materials in the number of expressed genes. Combined treatment such as ELICE16INDURES + BABA + SA at 0 day in comparison with control at day 2 showed large changes in gene expression (8 vs 9). This combined effect was experienced also at day 2 such as ELICE16INDURES + BABA and ELICE16INDURES + BABA + SA in comparison with control and SA at day 0 (15 vs 1 and 16 vs 1). SA treatment might strongly affect the gene expression numerically just the day 0 and 2 day as well (10 vs 1, 10 vs 2, 15 vs 2, 16 vs 2). Functional analyzes to determining genes that might be in the background of numerical data were performed (Figure 4B).

**Figure 4.**
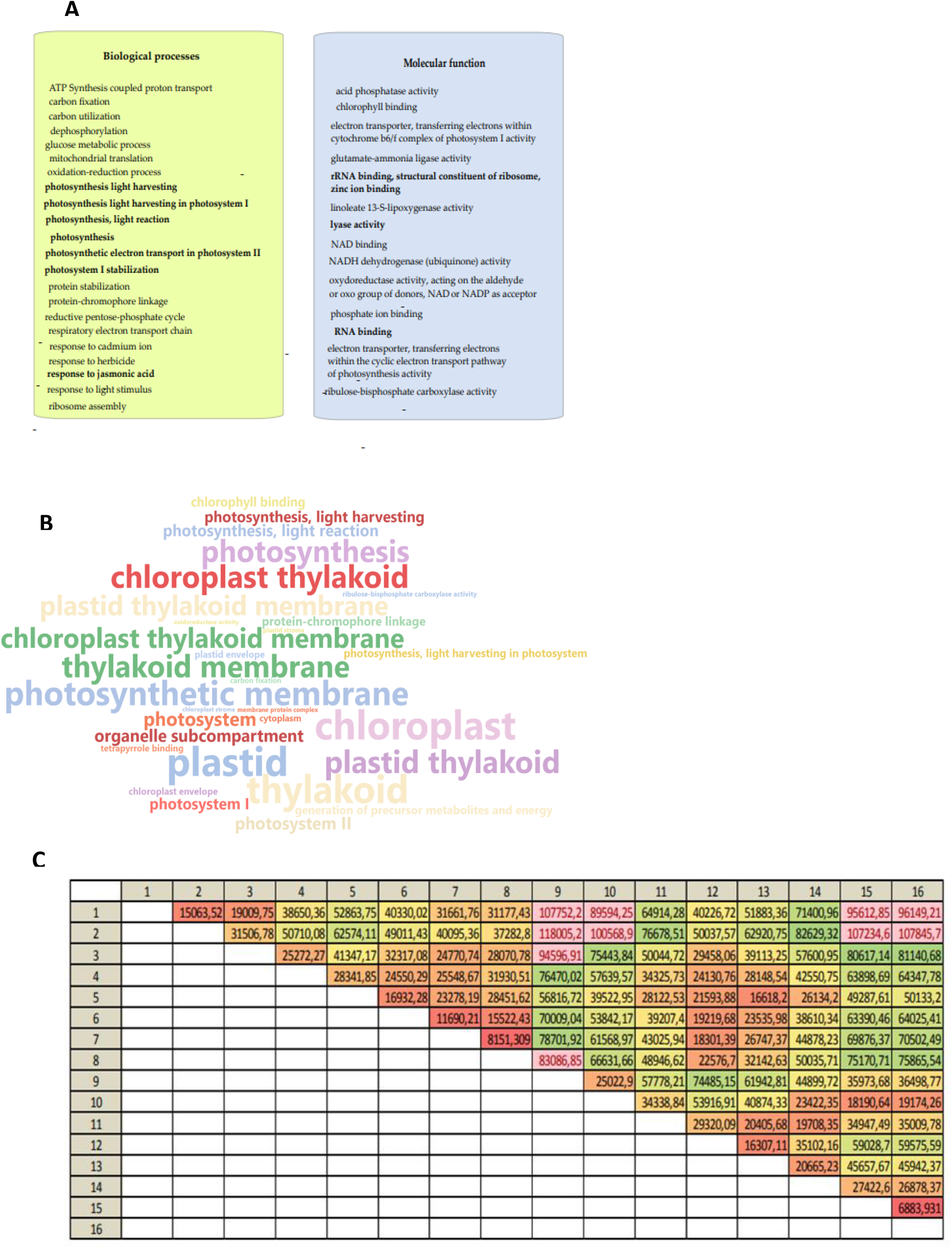
Searching of sample differences by 2-norm and determination of most variable genes by 1-norm Cell biochemical processes and molecular function of the top50 gene showing the largest change in the total data set of 16 samples determined by 1-norm are visualized **(A)**. Column combinations (16 × 15/2) in sample difference searching process by 2-norm **(C)**. Numbers means the treatments according to experimental design. Pink-colored values are the top 10 sample combinations with the highest expression differences. Columns showing the greatest differences for the total transcriptomic data may determine which two samples had the largest difference in gene expression at the total gene level; namely which 2 samples have the most different transcriptomic pattern. Values indicate the distances between the samples. A higher value means more different patterns. Sample pairs selected based on the table were used to perform pairwise gene expression analysis. To better understanding, a WordCloud **(B)** of gene set enrichment analysis of sample pairs screened with 2-norm was performed. Words mean GO categories of pink-colored values that were highlighted in **(C)**.

Sample pairs highlighted with pink showed a high photosynthetic activity associated with chloroplast thylakoid membrane, light-harvesting of photosystems I and II, without exception. This phenomenon showed a strong gene expression measured between days 0 and 2. Selected pairs such as (i) 1 vs 10 (ii) 1 vs 11; (iii) 1 vs 13; (iv) 1 vs 15; (v) 1 vs 16; and the outstanding (vi) 9 vs 8 combinations indicated also a strongly enhanced chloroplast activity. Since the enhanced photosynthetic processes were suppressed the other valuable gene expression data, pairwise data was analyzed always on the given day of treatments (Figure 5).

**Figure 5.**
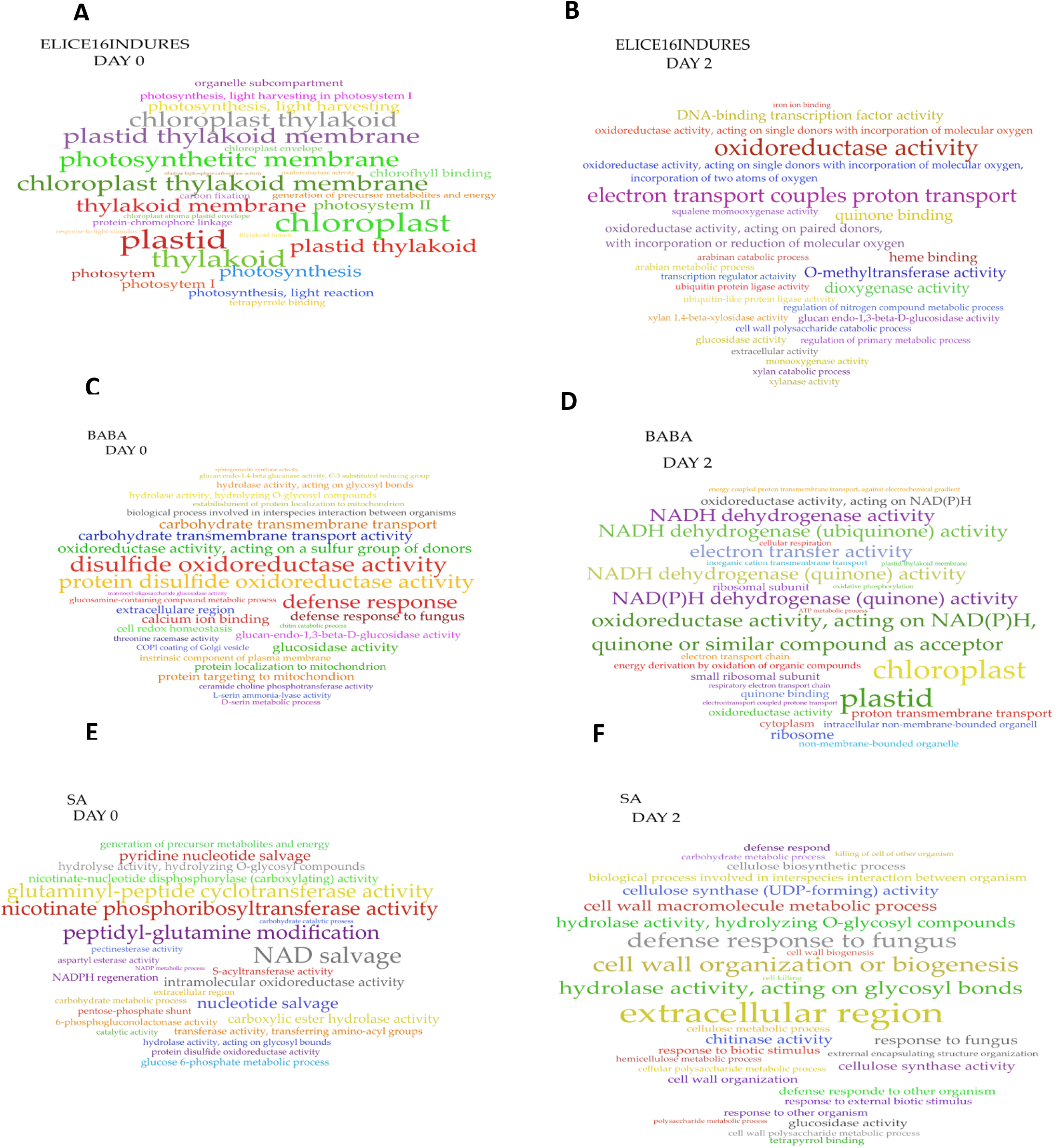
Functional annotation of pairwise differential expression of treatments. WordCloud representation of GO enrichment analysis of sample pairs. ELICE16INDURES vs Control **(A, B)**, BABA vs Control **(C, D)** and SA vs Control **(E, F)** were compared in time course at the days 0 and 2 and presented in the pairwise Clouds. ELICE16INDURES enhanced the photosynthesis just after addition, which effect was not observed analyzing SA

#### Determination of the most variable genes and their functions by 1-norm

Based on distance values according to 1-norm analysis, the top 50 genes showing the most significant changes across 16 treatments were determined and filtered out. These genes showed the highest changes among the samples. An annotation table was assigned to these records. Based on GO terms, we determined biochemical processes that were affected by the top50 genes. The biological process and molecular function of involved genes are summarized in Figure 4A. The screening shows that the genes with the most change in the entire data set are also involved in the biochemistry of the jasmonic acid pathway, which was observed in field experiments as well. Interestingly, pathways responsible for photosynthesis also showed a large change as a result of the treatments, which are consistent with the NDVI studies. The listed oxidation-reduction system, as well as the cadmium ion response associated with oxidative stress, suggested that the antioxidant enzyme system was also stimulated in the samples, which is consistent with previous literature data for salicylic acid and BABA applications (Guo et al., 2013; Pastor et al., 2013). From this study, we determined which biological processes are most affected by the treatments. The up-and down-regulation of these processes was in focus in our further experiments such as the jasmonic acid pathway, (ii) photosynthesis, (iii) cellular respiration, and (iv) regulation of oxidative stress.

#### Pairwise differential expression analysis

To help the selection of sample pairs that were analyzed, we based these comparisons on the differences determined by the 2-norm. Top50 DEGs of the pairs of single treatments on days 0 and 2 were determined using enrichment analysis (Fisher’s Exact Test). The DEGs were further performing GO analysis. The GO categories of over-expressed genes are illustrated using WordCloud based on GO IDs Fisher results (Figure 5A-F). These results indicated different mechanisms of action of the three investigated immune-priming agents. It can be seen that pathways of pathogen responses involved to priming are induced immediately after BABA addition which was associated with regulation of cellular redox homeostasis. Gene induction effect of BABA-priming decayed on day 2. In contrast, SA showed a prolonged effect on priming genes such as response to external biotic stress to fungus and other organisms was induced strongly on day 2. Genes involved in this process showed high hydrolase, glucosidase, or chitinase activities as described earlier for example in rice by pathogen-derived elicitors and SA (Silverman et al., 1995). We found that ELICE16INDURES enhanced the photosynthesis just after addition, which effect was not observed analyzing SA and BABA treated samples. Enhanced photosynthesis may explain the higher NDVI data observed in field experiments. Mechanism of action of this agent during priming indicated a strong redox activity and high expression of DNA binding transcription factors. The transcription factor regulation through redox changes associated to plant pathogen defense mechanisms were observed for just about 25 years (Wu et al., 1997; Zhang et al., 1999; Dangl and Jones, 2001; Mou et al., 2003; Wendehenne et al., 2014).

Transcription regulatory activity corresponded to the high expression of TIFY domain proteins. Further analysis of the ELICE16INDURES application on day 2 was performed making a heatmap and determine the top50 DEGs (Supplemental Figure S2). In this investigation sample 9 (Control of day 2) and sample 13 (ELICE16INDURES of day 2) were set up as test and reference data sets. Gene set enrichment analysis of top50 DEGs obtained from heatmap and GO information are summarized in Supplemental Table S3. We found overexpression of genes of TIFY 3A (TRINITY_DN489_c0_g1, TRINITY_DN966_c0_g1), TIFY 9 (TRINITY_DN354_c0_g1), TIFY 10A (TRINITY_DN8954_c0_g1), TIFY 11B (TRINITY_DN299_c0_g1) involving to JA response. *Arabidopsis* TIFY homologs, JAZ proteins act as negative regulators in JA signaling (Chini et al., 2007; Thines et al., 2007; Vanholme et al., 2007), and in response to JA and wounding TIFY domain proteins might be overexpressed. However, in cereal plants, TIFY mediated JA signal may work differently, hypothetically. In rice extensive up-regulation of OsTIFY10 and 11 members, moderated up-regulation of OsTIFY3 and OsTIFY9, and no change in the expression of OsTIFY1a and OsTIFY2a was observed in jasmonate associated growth. TIFY1a and TIFY2a were not differentially changed in our study as well. This was manifested by increased growth, grain weight, and flower number associated with shortened period to flowering. Since rice was not sensitive to external jasmonate addition, Hakata et al suggested that TIFY genes may promote growth by desensitizing plants to JA (Hakata et al., 2017). We observed yield increase in field conditions of ELICE16INDURES barley cultures corresponding to the findings of Hakata and al. Enhanced oxidoreductase activity suggested a strong effect of oxidative stress regulation of ELICE16INDURES which was just predicted during field experiments. JA response was mediated primarily by serine-threonine protein kinase signal transduction pathways. This may indicate the mechanism of action of ELICE16INDURES might be associated with receptor signaling of Jasmonate-mediated plant responses.

#### Determination of expression changes of genes of interest (GOIs)

Responses to certain or multiple stress factors are a complex task of plants in which central regulatory hormones such as SA, JA, ethylene, and ABA play key roles. Since the defense mechanisms are complex regulation of these hormones overlapping and linked – positive and negative -regulation may be observed (Ahmed et al., 2013). Many natural-based elicitors may induce a primed state in plants regulating hormonal pathways and modulating antioxidant systems (Aranega-Bou et al., 2014). Taking into account these regulatory pathways we investigated the gene expression changes in samples 1-16 in phytotron experiments. Genes associated with SA, JA, ABA pathways, and redox-system were analyzed *in silico*. Validation of *in silico* gene expressions was performed by RT-qPCR analyses (Figure 6 B-C).

**Figure 6.**
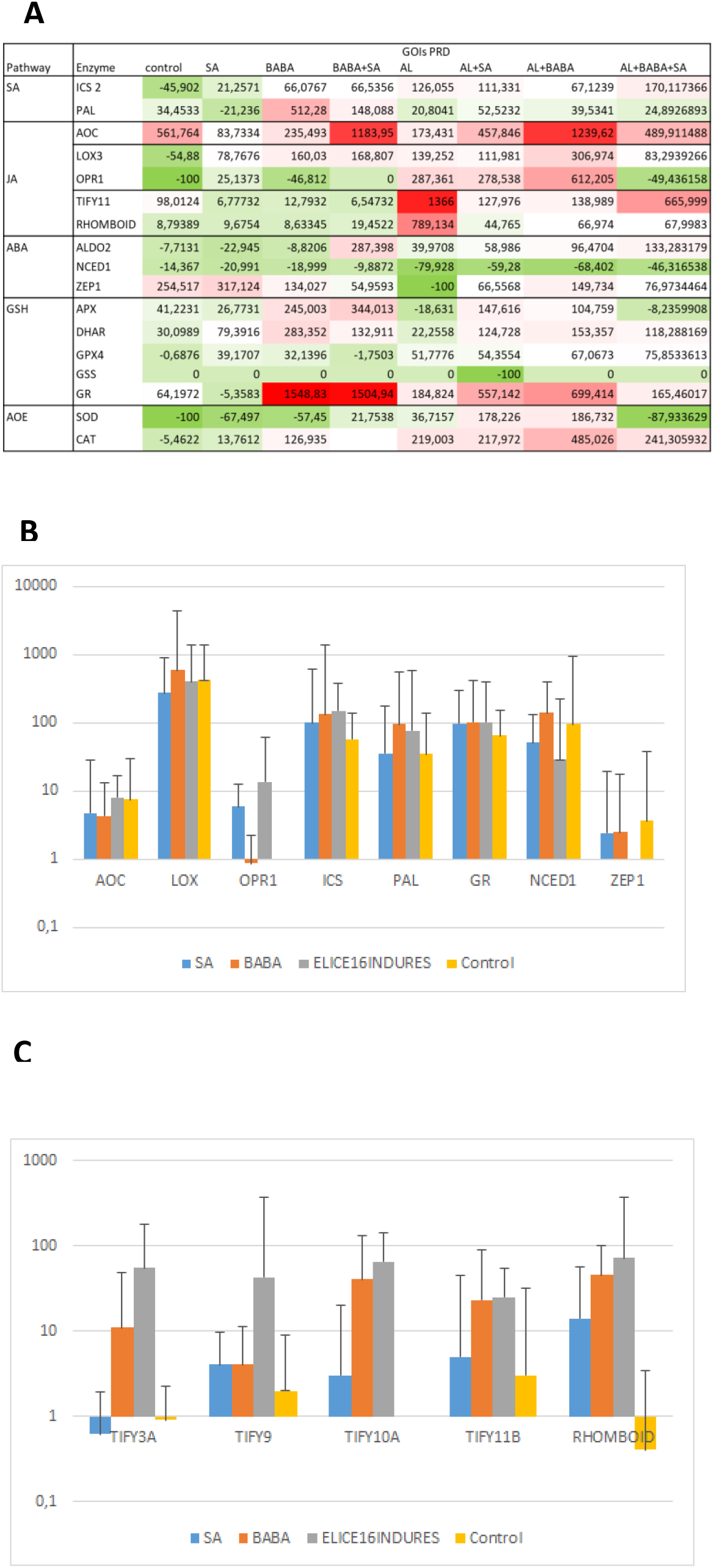
Validation analysis of genes of interest (GOI): ABA, JA, SA pathway-related genes, GR, TIFY-domain, and Rhomboid proteins. PRD values of genes **(A)**. Time course RPM changes as the response of treatments. Since expression intensity of each investigated gene showed high variances - in accordance with their physiological necessities - under control conditions, RPM deviation over days 0. and 2. were determined and expressed in %. Abbreviations: EL, ELICE16INDURES; SA, salicylic acid; BABA, beta-amino butyric acid; ABA, abscisic acid; JA, jasmonic acid; GSH, glutathione metabolism; AOE, antioxidant enzymes. RT-QPCR validation of SA, JA, ABA, GSH pathway genes treated by priming inducing materials **(B)**. RT-QPCR validation of TIFY domain proteins and RHOMBOID like 2 protein treated by priming inducing materials **(C)**. All of investigated priming active materials affected positively the expression of TIFY genes, however, it may be concluded that among them ELICE16INDURES induced most strongly these transcription factors. The deviations shown in the diagrams are linearly scaled at [min, max] intervals. Min: 0.06, max: 1.96 **(A)**; Min: 0.13 Max: 1.98 **(B)**

In or study 20 genes (GOIs) and their interconnected regulation were investigated involving SA, JA, ABA and AOS genes. Functional analysis of extracted cds was performed before mapping GEx reads. These annotations such as GO terms and enzyme pathway analysis are summarized in Supplemental Table S4. PRD was calculated to analyze how much gene expressions changed relative to themselves. Synergistic and antagonistic effects of exogenous addition of priming-elicitors SA and BABA were investigated and compared with ELICE16INDURES hypothetical priming-active compound.

##### Regulation of SA pathway genes

The key SA regulators are ICS and PAL which enzymes may help plants in their protection against environmental stresses and are modulated by different abiotic, biotic stress factors and during defense processes (Wildermuth et al., 2001). SA application can positively regulate these enzymes -such as high expression and enhanced activity -during drought, salt, chilling, heat, heavy metal and UV-B radiation stresses(Khan et al., 2015). In barley under drought conditions overexpression of ICS was correlated with a higher activity of antioxidant enzymes and lower levels of reactive oxygen species (Wang et al., 2021). Application of SA enhanced the activity of CAT, peroxidases, SOD, APX, GR under heavy metal stresses in different plant species (Arif et al., 2020). In our results, SA application upregulated ICS2 however repressed PAL by 20%. BABA and ELICE16INDURES treatments acted synergistically with exogenous SA addition to both genes. Correlation with GSH pathway and AOE genes were observed with exogenous SA application except for SOD that was repressed. Since during control conditions SOD was repressed as well, we supposed that SA treatment does not achieve its enhanced effect on the photosynthetic membrane and photosystem II activity involving the antioxidant system.

##### Regulation of JA pathway genes

Plant immunity especially involves JA biosynthesis and signaling in plant defenses against necrotrophic and insect infections (Brenya et al., 2020). During JA biosynthesis the chloroplast and the peroxisome are participating therefore lipoxygenase activity may play a key role in JA-induced priming. We detected enhanced LOX3 activity in all treatments suggesting a positive effect on the JA pathway. A similar effect was proved in the case of Vitamin B2 (Azami-Sardooei et al., 2010; Taheri and Tarighi, 2010). Furthermore, high AOC up-regulation was detected by BABA that could act synergistically with SA and ELICE16INDURES. Contemporarily BABA represses OPR3 in opposite with SA and ELICE16INDURES. Since OPR3 catalysis dnOPDSA to OPC6 step in peroxisomes, this suggests the effect of BABA may be concentrated in chloroplast-joint scenarios or Beta-oxidation in peroxisomes (Ruan et al., 2019). BABA showed a high effect acting antagonistically on the other two investigated materials. Based on the results of field experiments TIFY11 and RHOMBOID-like protein 2 were also analyzed in these samples. Time course positive regulation of TIFY11 and RHOMBOID-like protein 2 genes were observed in phytotron experiments in correspondence with field investigations. This phenomenon was detected in the case of ELICE16INDURES and combined treatment with BABA. However single BABA and SA treatments didn’t regulate these genes outstandingly. Transcriptional control of the JA pathway may be controlled by several factors (Caarls et al., 2015), therefore positive or negative gene regulation by different inducers is not surprising. Our results suggest that BABA and ELICE16INDURES induced outstanding changes in the JA-signaling, in contrast with SA which had less effect on these genes.

##### Regulation of ABA pathway genes

The accumulation of ABA is controlled by the enzyme 9-cis-epoxycarotenoid dioxygenase (NCED). Strong negative regulation of NCED1 by all treatments was found in this experiment. NCED1 was significantly down-regulated in drought stress, at which time ABA content reached a peak (Liu et al., 2016). However negative regulation of NCED1 is unclear and little studied. Our results indicated that plant responses to the investigated immune-priming activators might be manifested in the negative feedback of NCED1.

##### Regulation of GSH metabolism genes

The expression of antioxidant enzyme genes under different stresses was tested in a few studies on barley (Harb et al., 2015) which detail that abiotic and biotic stresses strongly induces expression of these genes promoting detoxification. We investigated how these genes may be altered to exogenous metabolic enhancers which may be associated with priming. BABA, SA and ELICE16INDURES synergistic and antagonistic effects on the regulation of GSH pathway and AOS were therefore analyzed. In plants, glutathione (GSH) metabolism is a key part of AOS and stress management. It is required for efficient defense against plant pathogens. Many enzymes use GSH as an electron donor or as a substrate during sulfur assimilation, flower development, salicylic acid, and plant defense signaling. We investigated the PRD values of GSH pathway key genes such as ascorbate peroxidase (APX), dehydroascorbate reductase (DHAR), GSH peroxidase (GPX), GSH reductase (GR) and GSH synthase (GSS).

GSS is the second enzyme in the GSH biosynthesis pathway. It catalysis the condensation of gamma-glutamylcysteine and glycine, to form glutathione. In *A. thaliana* low levels of GSS have resulted in increased vulnerability to stressors such as heavy metals and toxic organic chemicals (Maksymiec et al., 2005). The presence of a thiol functional group allows its product GSH to serve both as an effective oxidizing and reducing agent in numerous biological scenarios (Saez et al., 1990). GSS was changed negatively only in the case of SA treatment, other treatments did not affect this gene.

In plants, GR catalyzes the reduction of GSH disulfide (GSSG) to the sulfhydryl form GSH, which is a critical molecule in resisting oxidative stress and maintaining the reducing environment of the cell. It is part of the nglutathione-ascorbate cycle in which reduced GSH reduces dehydroascorbate, a reactive byproduct of the reduction of hydrogen peroxide. In particular, GR contributes to plants’ response to abiotic stress. The activity of enzymes was modulated in response to metals, metalloids, salinity, drought, UV radiation, and heat-induced stress (Gill et al., 2013). A strong up-regulation in GR expression was detected after BABA treatment and down-regulation after SA treatment. Exogenous addition of SA and ELICE16INDURES may impair the BABA effect. However, the synergistic effect on GR up-regulation was observed in the case of SA + ELICE16INDURES combination.

GPXs enzyme family possesses nine stress-related genes with conserved domain and high homology in plant species. These enzymes reduce lipid hydroperoxides to their corresponding alcohols and reduce free hydrogen peroxide to water. GPXs are induced by oxidative stress (Khan et al., 2020) pathogen infections (Pieczul et al., 2020), mechanical stimulation (Singh et al., 2020), salt, cold, drought and metal treatments (Han et al., 2020).

DHAR regulates the cellular ascorbic acid redox state, which in turn affects cell responsiveness and tolerance to environmental ROS. DHAR is important for plant growth such as maintain steady-state chlorophyll and Rubisco (Chen and Gallie, 2006). Upregulation of GPX and DHAR was observed in all treatments except ELICE16INDURES.

APX is part of the ascorbate–glutathione cycle, which requires ascorbate to scavenge H_2_O_2_ (Reddy et al., 2004; Sousa et al., 2015). APX was regulated negatively by ELICE16INDURES which effect may be slightly observed in the combined treatments (ELICE16INDURES + BABA + SA, ELICE16INDURES + BABA). A synergistic effect on the APX up-regulation was observed in BABA + SA combined treatment.

To summarize, GSH metabolism may be accelerated by the investigated priming-active substances, among which the most effective was the BABA treatment. This effect is outstanding in the case of GR. The other two substances show higher activity in combination suggesting synergistic effects.

##### Validation of enhanced expression of TIFY family members and RHOMBOID like protein 2

The role of TIFY-motif proteins in JA signaling is known (Chung and Howe, 2009) however which stress factors may activate the TIFY-mediated events is less studied. In this study applying a comprehensive genome-wide bioinformatics analysis, we experienced that JA response was strongly associated with thylakoid membrane scenarios at the gene expression level. This phenomenon was remarkable triggering priming-associated genes by ELICE16INDURES. The expression profile performed by *in silico* predictions was validated by the RT-qPCR technique using plants cultivated in the phytotron. TIFY3A, TIFY9, TIFY10A, TIFY11B and RHOMBOID-like protein 2 showed a strong up-regulation after ELICE16INDURES treatment at day 2. Expression of these genes was observed after BABA and SA treatment at a significantly lower rate. TIFY3A was downregulated by SA. All substances triggered the relative expression of the RHOMBOID protein. Since rhomboid proteins are proved to be associated with the internal region of the thylakoid membrane we concluded that ELICE16INDURES possess a priming effect through JA signaling. This hormonal regulation takes place in the chloroplast. Since we observed augmented photosynthetic areas in field experiments meaning an increased amount of chloroplasts in ELICE16INDURES treated plots, we concluded that higher gene levels of JA pathway genes (AOC, LOX, OPR1) correlated also with enhanced TIFY and Rhomboid proteins in phytotron experimental setup (Figure 6 A-C). The role of chloroplast-Rhomboid proteases was studied in *A. thaliana* indicating to have a positive impact on fertility and flowering through JA signaling (Adam, 2015)

Plant immune response may be interconnected by various metabolic pathways, among which the most important is the photosynthetic activity from farming aspects. Alterations in physiological chloroplast events lead to plant wilting therefore understanding chloroplast function associated with priming is considerable. Since chloroplast is a major synthesis site for many plant hormones such as ABA, JA, SA, and free radical production, there is a complex relationship between photosynthesis and defense-related signals (Lu and Yao, 2018) which scenarios take place in thylakoid membranes, primarily. The rhomboid family of serine proteases plays a pivotal role in a diverse range of pathways, activating and releasing proteins via thylakoid intramembranous proteolysis. The plastid intramembranous proteolysis of RHOMBOIDs was studied in *A. thaliana* and found that the lack of chloroplast-located rhomboid proteases caused reduced fertility and aberrations in flower morphology which was interconnected to JA signaling (Adam, 2015).

## MATERIALS AND METHODS

### Priming-active compounds

ELICE16INDURES was produced in the Institute of Medicinal Plant and Herbs, Ltd, Hungary. We used the high-pressure extraction with supercritical carbon dioxide as a solvent (scCO_2_ extraction). 11 scCO_2_ extracts of medicinal plants were establishes in a common set of states and encapsulated in sunflower lecithin based liposomes (Figure 1 D). The ratio of scCO_2_ extracts see in Figure 1B. We reach to entrap the active agents into small multilamellar vesicles (MLV) of 250-350 nm by using active trapping techniques (Mayer et al., 1986). Transmission electron microscopy (TEM) was used to characterize the sizes and structures of liposomes in the range 10-1000 nm. Liposome structures were visualized and analyzed by TEM (Figure 1). Active compounds of ELICE16INDURES are summarized in Supplemental Table S5. scCO_2_ extracts were purchased from FLAVEX Naturextrakte GmbH, Germany. The two immune priming activator compounds SA and BABA (purity 99 and 97%) were purchased from Merck (MilliporeSigma, US).

### TEM processing and particle size analysis

Samples for transmission electron microscopy (TEM) were prepared by diluting liposome concentrates with MQ water and depositing a drop of the suspension on copper TEM grids covered by continuous carbon amorphous support film. After 5 minutes the drop was blotted with filter paper. Diluted (2%) phosphotungstic acid was used for sample staining. The staining time was 30 seconds. Measurement conditions were as follows: TEM analyses were performed using a Talos F200X G2 instrument (Thermo Fisher), operated at 200 kV accelerating voltage, equipped with a field-emission gun and a 4096×4096 CMOS camera for imaging. In our study TEM bright-field images were collected at 36000x magnification for particle size analysis and other magnifications as well to visualize the details and the inner structures of the liposomes.

Particle size analysis was performed based on TEM bright-field images using Gimp and ImageJ software. The Gimp software was used for image processing (including contrast enhancement, blurring, and manual outlining) and ImageJ was used for creating the final image with grayscale threshold settings and for an analysis of particle sizes. The smaller particles had continuous borders and sufficient contrast for distinguishing them from their background with proper threshold settings, enabling ImageJ to recognize these particles. We could not use this method directly on the larger particles because of their diffuse details or discontinuous borders. Therefore, we manually outlined and filled the larger particles with black color to create an image with white background and black particles. As the last step, we removed the black pixels forming bridges between the particles to create the final image for the particle size analysis.

### Cultivation of plant materials and chemical treatment

Fresh leaves were collected on days 0 and 2 after treatments from 11-16-day-old autumn barley plants (diploid). Day 0 means 2-3 hours after treatment. Plants were cultivated in phytotron and arable fields.

#### Phytotron experiments

Exogenous treatment of SA, BABA, and ELICE16INDURES was performed in phytotron experiments. Plant growth chamber type was MLR352HPA -115V NEMA 5-20, 220V / 60Hz – Panasonic. Treatment conditions were as follows: temperature during the 1. day and night was 25 °C. The temperature during the 2-16. days and nights were 25 °C and 15 °C. Duration of the day was 12 hours, 04-4 p.m. Treatments were as follows: Na-SA (MW: 160.11 g/mol), 300 µM-solution; BABA (MW: 103.121 g/mol), final concentration in soil was 25 µM; ELICE16INDURES, 1 ml/100ml water. Experimental design and sample collection were performed according to Figure 3A and B.

#### Field experiments

Plants in field experiments were sprayed by TTAM4E drone with low and high doses of ELICE16INDURES. We used a positive control that was a commercially available plant conditioner, Fitokondi^®^. Applied doses and plot allocations were: 1, control without treatment; 2, positive control treated with Fitokondi; 3, 10 g/ha; 4, 20 g/ha; 5, 30 g/ha; 6, 60 g/ha; 7, 120 g/ha ; 8, 240 g/ha ELICE16INDURES. Plot allocation of doses see in Figure 2A. After sample collections, samples were prepared for RNA-sequencing and gene expression profiling. Three biological repeats were investigated.

### Determination of normalized difference vegetation index (NDVI) by remote sensing

To monitor NDVI in the field population we used a DJI-phantom 4 agro drone equipped with a near-infrared camera. The recording was carried out on day 2 after treatment. Single aerial pictures were combined afterward with AgiSoft Photoscan Professional software using high-quality dense cloud processing and mesh construction settings (Figure 2A). To NDVI calculation after identifying the plots based on combined aerial photographs, sample areas were cut out using self-developed software. Sample collection for molecular biological analysis was performed at the same day.

### Preparation of RNA-seq libraries

Approximately 30 mg of plant tissues were placed in a 1.5 ml Eppendorf LoBind tube containing glass beads (1.7-2.1 mm diameter, Carl Roth, Karlsruhe, Germany) and 100 µl of TRI-Reagent (Zymo Research). The Eppendorf tube was firmly attached to a SILAMAT S5 vibrator (Ivoclar Vivadent, Schaan, Liechtenstein) to disrupt and homogenize the tissue for 2×15s. Total RNA was extracted using Direct-zol™ RNA MiniPrep System (Zymo Research) according to the manufacturer’s protocol. The RNA Integrity Numbers and RNA concentration were determined by RNA ScreenTape system with 2200 Tapestation (Agilent Technologies, Santa Clara, CA, USA) and RNA HS Assay Kit with Qubit 3.0 Fluorometer (Thermo Fisher Scientific, Waltham, MA, USA), respectively. For Gene Expression Profiling (GEx) library construction, QuantSeq 3’ mRNA-Seq Library Prep Kit FWD for Illumina (Lexogen GmbH, Wien, Austria) was applied according to the manufacturer’s protocol. The quality and quantity of the library were determined by using High Sensitivity DNA1000 ScreenTape system with 2200 Tapestation (Agilent Technologies, Santa Clara, CA, USA) and dsDNA HS Assay Kit with Qubit 3.0 Fluorometer (Thermo Fisher Scientific, Waltham, MA, USA), respectively. Pooled libraries were diluted to 1.8 pM for 1×86 bp single-end sequencing with 75-cycle High Output v2 Kit on the NextSeq 550 Sequencing System (Illumina, San Diego, CA, USA) according to the manufacturer’s protocol. By using longer reads QuantSeq FWD allows to exactly pinpoint the 3’ end of poly(A) RNA and therefore obtain accurate information about the 3’ UTR. Using this sequencing fragments of coding sequences are 260-300 bp long on average.

### Bioinformatics analysis – reads processing

3 × 16 libraries were sequenced with a final output single-end, 15-18 M x 80 bases long. Quality control (QC), trimming, and filtering of .fastq files were performed in preprocessing step. The QC analysis was performed with FastQC (Andrews et al., 2010) software. For all the 3 × 16 libraries the Phred-like quality scores (Qscores) were set to >30. Poor quality reads, adapters at the ends of reads, limited skewing at the ends of reads were eliminated by using Trimmomatic (Bolger et al., 2014). Contamination sequences and N’s were filtered out with a self-developed application GenoUtils as described earlier (Mátyás et al., 2019): reads containing N’s more than 30 % were eliminated; reads with lower N’ ratio were trimmed with a final length > 65. Reads passed of preprocessing were further assembled and analyzed.

### *De novo* assembly and mapping

Reference transcript dataset was firstly created from the libraries (TrinityH16-transcript, Supplemetal File S1). To increase the coverage of the transcripts *de novo* assembly was performed with all the cleaned reads of combined 3 × 16 libraries. To obtain longer mRNA fragments from the combined short reads (65-80bp) without reference Trinity assembler with 23K-mer were used (Grabherr et al., 2011). For *de novo* assembly and mapping we used a server with 512 GB (Gigabytes) of RAM, 64 cores (CPUs), and Ubuntu as the operating system. To assess the read composition of the assembly, input RNA-Seq reads were aligned to the transcriptome assembly using Bowtie2 (Langmead and Salzberg, 2012). All reads from each biological repeat (3 × 16) were separately mapped against the reference dataset. Reads mapped to the assembled transcript were captured. Statistic of mapped reads was performed and compared between all replicates. We used TrinityH16-transcript for gene expression profiling. Collapsing of splicing isoforms were performed with OmicsBox and SuperTranscripts (gene-level assembly) were used in further investigations TrinityH16-supertranscript, Supplemental Data Set S2. Trinity H-field (Supplemental Data Set S6) transcript dataset of filed experiments were performed with the same method.

### Gene level quantification

To estimate gene expression from RNA-sequencing CountTable was created (Supplemental Data Set S3). To count how many reads map to each feature of interest (transcripts) each sample reads were aligned to the reference TrinityH16-supertranscript. Count Table creation was performed with omixbox.biobam (OmicsBox, https://www.biobam.com/omicsbox) using the HTseq package (Anders et al., 2015). Based on the data of CountTable further analyses were performed such as differential expression analysis and mathematical prediction of the test area.

### Pairwise differential expression analysis

Numerical analysis of differentially expressed genes (DEGs) in a pairwise comparison of two different experimental conditions -*gene expression analysis* – was carried out using omixbox.biobam (OmicsBox, https://www.biobam.com/omicsbox). The used application is based on the edgeR program implementing quantitative statistical methods to evaluate the significance of individual genes between two experimental conditions (Robinson et al., 2010). TMM (Weighted trimmed mean of M-values) normalization method was performed.

### Functional annotation

Functional annotation and Gene Ontology (GO) analysis were carried out using OmixBox.Biobam as follows: 1. Sequences were blasted against NCBI nr (non-redundant) Viridiplantae database (downloaded in 2019) applying blastn configuration locally; 2. To retrieving GO terms associated with the 10 Hits obtained by the Blast search GO mapping and annotation were performed. GeneBank identifiers (gi), the primary blast Hit ids, were used to retrieve UniProt IDs making use of a mapping file from PIR (Non-redundant Reference Protein Database) including PSD, UniProt, Swiss-Prot, TrEMBL, RefSeq, GenPept and PDB. Accessions were searched directly in the dbxref table of the GO database. BLAST result accessions were searched directly in the gene-product table of the GO database; 3. GO annotations were specified according to GO terms: molecular function, cellular component, biological process.

### Determination of uniquely expressed transcripts

Count table data of each biological replicates were used. According to the mapping statistics, there were no significant differences between the biological replicates. Average values of counts were taken into account used to create a 60,615 (number of genes, rows) x 16 (number of samples, column) matrix from the count table. Normalization of mapped read counts, RPM index (reads per million mapped reads, Figure 7A) were firstly calculated from matrix data for the determination of uniquely expressed transcripts and mathematical modeling of interest area (see below). Highly characteristic transcripts that were uniquely expressed were filtered out from the matrix with Microsoft access database manager. Identifiers of these transcripts in each treatment condition (sample) were separated and saved in txt files. To the better evaluation graphical representation of results was grouped into 4 Venn diagrams (Figure 3 C-F) using the web-based interactivenn.net (Heberle et al., 2015).

**Figure 7.**
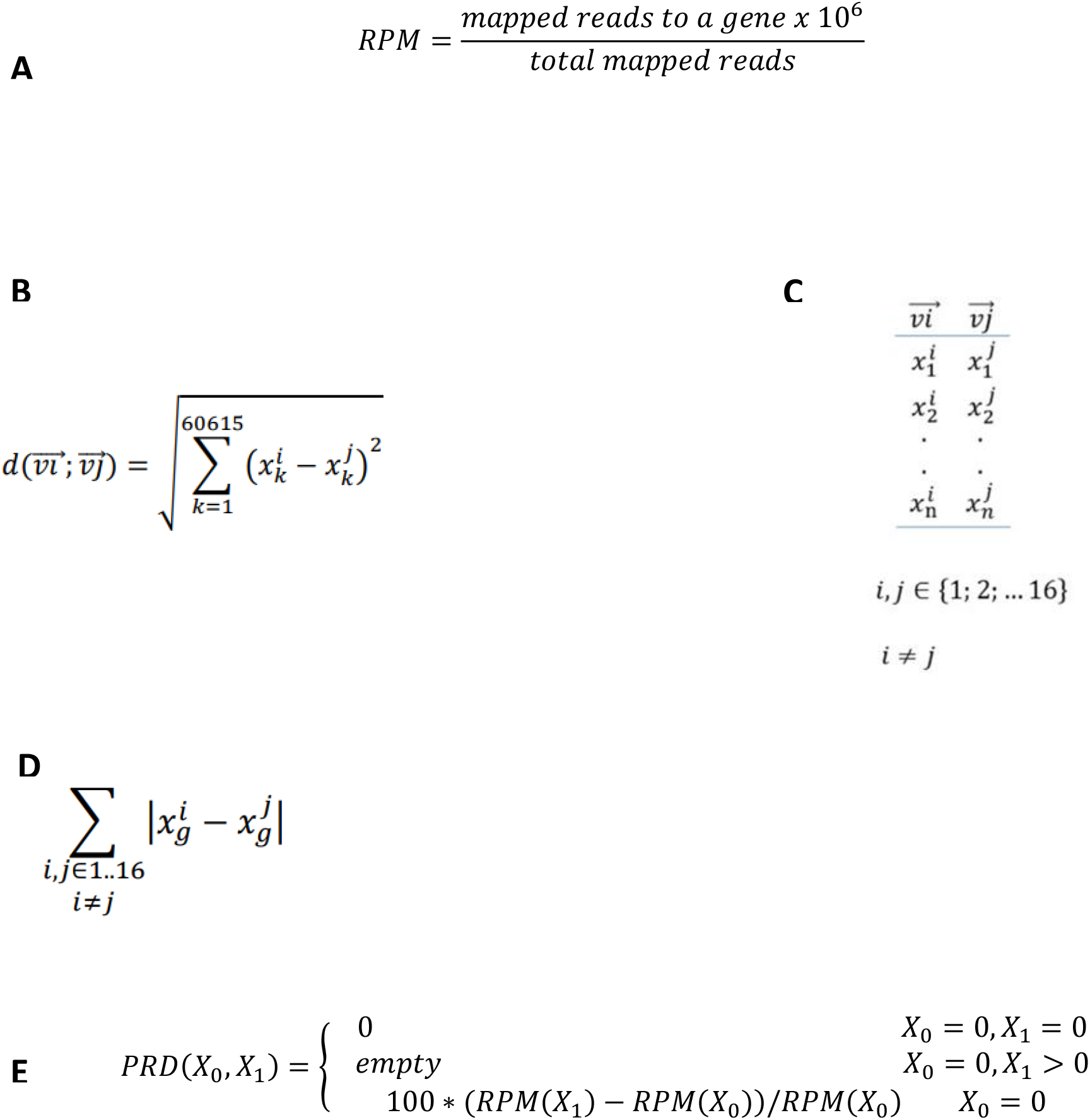
Formules used for calculations. **A**. Equation of RPM index used to screening uniquely expressed transcripts and mathematical modeling of interest area. **B**. Equation of distance calculation according to the 2-norm. **C**. 2 elements of the examined n-dimensional vector space, where n = 60,615, ***vi*** and ***vj*** vectors are column vectors of the 1-16 samples in the count table **D**. Equation of change difference of a transcript between the sample pairs. Symbols: **g** is the given transcript of the gene; **xg** is the RPM value that belongs to the transcript; **i**,**j** are indexes of the sample pairs. **E**. Percentage of relative difference measured by in investigated genes (GOIs). **X0:** RPM value on day zero, **X1:** RPM value on day second, PRD: percentage relative difference

### Enrichment analysis

To determine cellular responses and cell-biochemical processes altered between 2 samples and how they are affected by the treatments, Fisher’s Exact Test was used. Fisher’s Exact Test performs functional analysis of differential expressed genes working with the entire annotated data set previously determined (Huang et al., 2009).

### Mathematical prediction of investigated area examination

To predict which treatment conditions show the most different transcriptomic events, search and determination the extremal points of the 16 transcriptome data sets as a multidimensional space. The 60,615 × 16 matrix was analyzed by column and by rows which fields contained the RPM indexes.

### Searching of sample differences by 2-norm

2-norm is a functional on vector space that assigns a non-negative (real) number to the elements of vector space. Hölder norms (e.g., 1-norm, 2-norm, p-norm) can be used to manage distances (length) in abstract spaces. In our case, we took a 60,615-dimensional vector space (with its 16 vectors, Figure 7C), in which the column vectors were spaced. We wondered how far apart these vectors in pairs are. Since the distance function is commutative, therefore in a given comparison, only the two different vectors were counted, their order was not. The examination number was given by a combination of 16 elements without second class repetition which is 16 * 15/2. The distance of 2 columns was given by the value of the vector of the difference between the two vectors according to norm 2 (Figure 7 B, C).

### Searching of differences between genes by 1-norm

During this method transcripts showing the largest changes in RPM across the 16 samples were determined. For this analysis, all 60,615 transcripts were involved. For each transcripts, absolute values of the difference in RPM values between the sample pairs were calculated and these values were summarized. We assumed that higher values are stronger in expression. As a result of the analysis, the top 50 genes were determined (Figure 7D).

### Determination of the most variable cell-biochemical pathways

The top 50 transcripts based on distance values by 1-norm were categorized according to GO terms. Cell biochemical processes were examined in more detail. Changes in cell responses as a result of treatment were grouped based on enrichment analysis (Huang et al., 2009).

### Determination of expression changes of genes of interest (GOIs)

We identified the accurate coding sequences (cds) of investigated pathway genes in our plant material. Therefore we used a transcriptome dataset from deep sequencing of three, treated and control phytotron samples of *H. vulgare* (TrinityH-Deep, Supplemental Data Set S4). Sequencing protocol and assembly were used as described earlier (Virág et al., 2016). We identified cds of 20 genes of interest (Supplemental Data Set S5). JA, ABA, SA and GSH pathway genes were determined. Contigs of these genes were identified based on *Oriza sativa* and *H. vulgare* using BLASTn with an E value less than 10^−5^ (Altschul et al., 1990). ORF (open reading frame) was determined with NCBI ORF finder. These cds were used as reference sequences to mapping reads of GEx libraries to determine gene expression changes. Time course RPM changes (PRD, percentage of relative difference) as a response to treatments were calculated (formula see in Figure 7E). Expression changes were calculated mapping reads of QuantSeq 3’ mRNA-Seq libraries in which one fragment per transcrinpt was produced, therefore, no length normalization was required. This method allows a more accurate determination of expression values in gene expression studies.

### Gene expression analysis with RT-qPCR amplification

RT-qPCR experiment of GOIs was performed as described earlier (Mátyás et al., 2019). Primers were designed with Primer3 (Koressaar and Remm, 2007; Untergasser et al., 2012) based on *in silico* sequences of GEx libraries. Gene expression analysis was established based on three technical and biological replicates and normalized with the reference gene glyceraldehyde-3-phosphate dehydrogenase, *GAPDH*.

### Accession Numbers

Raw reads of this project used for field and phytotron experiments (21 SRA experiments) are deposited in the NCBI SRA database under the accessions: Bioproject, PRJNA721578; Deep RNA sequencing, SRX10603947-SRX10603949; Sequencing of field experiments, SRX10598683, SRX10598684; Sequencing of phytotron experiments, SRX10600133-SRX10600148.

## Supplemental data

**Supplemental Table S1** Annotation of most interesting genes from the top50 DEGs in field experiments.

**Supplemental Table S2** Results of functional annotation of TrinityH16-supertranscript – reference-transcript dataset.

**Supplemental Table S3** Annotation of top50 DEGs of pairwise comparison

**Supplemental Table S4** Gene ontology and enzyme annotation of investigated sequences of SA, JA, ABA and GSH pathway genes.

**Supplemental Table S5** Biologically active components of ELICE16INDURES.

**Supplemental Figure S1** The top20 GO term categories of the reference transcriptome

**Supplemental Figure S2** Heatmap of the investigated 16 H. vulgare samples

**Supplemental Data Set S1** TrinityH16-transcript.fasta

**Supplemental Data Set S2** TrinityH16-supertranscript.fasta

**Supplemental Data Set S3** Count Table

**Supplemental Data Set S4** TrinityH-Deep

**Supplemental Data Set S5** Coding sequences and primers GOIs

## ACKNOWLEDGMENTS

The work was supported by the KFI_16-1-2017-0457 -*Development and production of a plant-based pesticide-plant conditioner for use in organic farming* -project of the Hungarian Government. We express our thanks to the Xenovea Ltd, Hungary to perform NGS library preparation and sequencing. We are grateful to the editor and reviewers for the valuable comments that have helped to improve the manuscript.

## AUTHOR CONTRIBUTIONS

E.V., and J.P.P., designed the project; G.H., M.K., B.L., L.G., Á.N., P.P., Z.T., performed experiments; K.D., B.K., K. S., E.V., and G.H., analyzed the data., E.V., and G.H., performed bioinformatics analyses., E.V., and Á.J., wrote the paper.

## REFERENCES

Abdel-Aziz, H., Hasaneen, M., and Omer, A. (2019). Impact of engineered nanomaterials either alone or loaded with NPK on growth and productivity of French bean plants: Seed priming vs foliar application. South African Journal of Botany 125, 102–108.

Adam, Z. (2015). Plastid intramembrane proteolysis. Biochimica et Biophysica Acta (BBA)-Bioenergetics 1847, 910–914.

Adeniyi, S., Orjiekwe, C., Ehiagbonare, J., and Arimah, B. (2010). Preliminary phytochemical analysis and insecticidal activity of ethanolic extracts of four tropical plants (Vernonia amygdalina, Sida acuta, Ocimum gratissimum and Telfaria occidentalis) against beans weevil (Acanthscelides obtectus). International Journal of Physical Sciences 5, 753–762.

Ahmed, I.M., Cao, F., Zhang, M., Chen, X., Zhang, G., and Wu, F. (2013). Difference in yield and physiological features in response to drought and salinity combined stress during anthesis in Tibetan wild and cultivated barleys. PloS one 8, e77869.

Altschul, S.F., Gish, W., Miller, W., Myers, E.W., and Lipman, D.J. (1990). Basic local alignment search tool. Journal of molecular biology 215, 403–410.

Anders, S., Pyl, P.T., and Huber, W. (2015). HTSeq—a Python framework to work with high-throughput sequencing data. bioinformatics 31, 166–169.

Andrews, S., Krueger, F., Seconds-Pichon, A., Biggins, F., and Wingett, S. (2010). FastQC: a quality control tool for high throughput sequence data. Babraham Bioinformatics. 2010.

Aranega-Bou, P., de la O Leyva, M., Finiti, I., García-Agustín, P., and González-Bosch, C. (2014). Priming of plant resistance by natural compounds. Hexanoic acid as a model. Frontiers in plant science 5, 488.

Arif, Y., Sami, F., Siddiqui, H., Bajguz, A., and Hayat, S. (2020). Salicylic acid in relation to other phytohormones in plant: a study towards physiology and signal transduction under challenging environment. Environmental and Experimental Botany, 104040.

Azami-Sardooei, Z., França, S.C., De Vleesschauwer, D., and Höfte, M. (2010). Riboflavin induces resistance against Botrytis cinerea in bean, but not in tomato, by priming for a hydrogen peroxide-fueled resistance response. Physiological and Molecular Plant Pathology 75, 23–29.

Beckers, G.J., and Conrath, U. (2007). Priming for stress resistance: from the lab to the field. Current opinion in plant biology 10, 425–431.

Bolger, A.M., Lohse, M., and Usadel, B. (2014). Trimmomatic: a flexible trimmer for Illumina sequence data. Bioinformatics 30, 2114–2120.

Brenya, E., Chen, Z.-H., Tissue, D., Papanicolaou, A., and Cazzonelli, C.I. (2020). Prior exposure of Arabidopsis seedlings to mechanical stress heightens jasmonic acid-mediated defense against necrotrophic pathogens. BMC Plant Biology 20, 1–16.

Burketova, L., Trda, L., Ott, P.G., and Valentova, O. (2015). Bio-based resistance inducers for sustainable plant protection against pathogens. Biotechnology advances 33, 994–1004.

Caarls, L., Pieterse, C.M., and Van Wees, S. (2015). How salicylic acid takes transcriptional control over jasmonic acid signaling. Frontiers in plant science 6, 170.

Calvo, P., Nelson, L., and Kloepper, J.W. (2014). Agricultural uses of plant biostimulants. Plant and soil 383, 3–41.

Chen, Z., and Gallie, D.R. (2006). Dehydroascorbate reductase affects leaf growth, development, and function. Plant Physiology 142, 775–787.

Chini, A., Fonseca, S., Fernandez, G., Adie, B., Chico, J., Lorenzo, O., García-Casado, G., López-Vidriero, I., Lozano, F., and Ponce, M. (2007). The JAZ family of repressors is the missing link in jasmonate signalling. Nature 448, 666–671.

Chung, H.S., and Howe, G.A. (2009). A critical role for the TIFY motif in repression of jasmonate signaling by a stabilized splice variant of the JASMONATE ZIM-domain protein JAZ10 in Arabidopsis. The Plant Cell 21, 131–145.

Dangl, J.L., and Jones, J.D. (2001). Plant pathogens and integrated defence responses to infection. nature 411, 826–833.

Divya, B., Suman, B., Venkataswamy, M., and Thyagaraju, K. (2017). A study on phytochemicals, functional groups and mineral composition of Allium sativum (garlic) cloves. International Journal of Current Pharmaceutical Research 9, 42–45.

Du Jardin, P. (2015). Plant biostimulants: definition, concept, main categories and regulation. Scientia Horticulturae 196, 3–14.

Dutta, P. (2018). Seed priming: new vistas and contemporary perspectives. In Advances in seed priming (Springer), pp. 3–22.

Farooq, M., Usman, M., Nadeem, F., ur Rehman, H., Wahid, A., Basra, S.M., and Siddique, K.H. (2019). Seed priming in field crops: potential benefits, adoption and challenges. Crop and Pasture Science 70, 731–771.

Figueiredo, A., Monteiro, F., and Sebastiana, M. (2015). First clues on a jasmonic acid role in grapevine resistance against the biotrophic fungus Plasmopara viticola. European Journal of Plant Pathology 142, 645–652.

Geelen, D., and Xu, L. (2020). The Chemical Biology of Plant Biostimulants. (John Wiley & Sons, Incorporated).

Gill, S.S., Anjum, N.A., Hasanuzzaman, M., Gill, R., Trivedi, D.K., Ahmad, I., Pereira, E., and Tuteja, N. (2013). Glutathione and glutathione reductase: a boon in disguise for plant abiotic stress defense operations. Plant Physiology and Biochemistry 70, 204–212.

Grabherr, M.G., Haas, B.J., Yassour, M., Levin, J.Z., Thompson, D.A., Amit, I., Adiconis, X., Fan, L., Raychowdhury, R., and Zeng, Q. (2011). Trinity: reconstructing a full-length transcriptome without a genome from RNA-Seq data. Nature biotechnology 29, 644.

Guo, Q., Meng, L., Mao, P.-C., Jia, Y.-Q., and Shi, Y.-J. (2013). Role of exogenous salicylic acid in alleviating cadmium-induced toxicity in Kentucky bluegrass. Biochemical Systematics and Ecology 50, 269–276.

Hakata, M., Muramatsu, M., Nakamura, H., Hara, N., Kishimoto, M., Iida-Okada, K., Kajikawa, M., Imai-Toki, N., Toki, S., and Nagamura, Y. (2017). Overexpression of TIFY genes promotes plant growth in rice through jasmonate signaling. Bioscience, biotechnology, and biochemistry 81, 906–913.

Han, L.-m., Hua, W.-p., Cao, X.-y., Yan, J.-a., Chen, C., and Wang, Z.-z. (2020). Genome-wide identification and expression analysis of the superoxide dismutase (SOD) gene family in Salvia miltiorrhiza. Gene 742, 144603.

Harb, A., Awad, D., and Samarah, N. (2015). Gene expression and activity of antioxidant enzymes in barley (Hordeum vulgare L.) under controlled severe drought. Journal of Plant Interactions 10, 109–116.

Heberle, H., Meirelles, G.V., da Silva, F.R., Telles, G.P., and Minghim, R. (2015). InteractiVenn: a web-based tool for the analysis of sets through Venn diagrams. BMC bioinformatics 16, 1–7.

Huang, D.W., Sherman, B.T., and Lempicki, R.A. (2009). Bioinformatics enrichment tools: paths toward the comprehensive functional analysis of large gene lists. Nucleic acids research 37, 1–13.

Kalbi, S., Fallah, A., and Shataee, S. (2014). Estimation of forest attributes in the Hyrcanian forests, comparison of advanced space-borne thermal emission and reflection radiometer and satellite poure I’observation de la terre-high resolution grounding data by multiple linear, and classification and regression tree regression models. Journal of Applied Remote Sensing 8, 083632.

Kanaoka, M.M., Urban, S., Freeman, M., and Okada, K. (2005). An Arabidopsis Rhomboid homolog is an intramembrane protease in plants. FEBS letters 579, 5723–5728.

Karny, A., Zinger, A., Kajal, A., Shainsky-Roitman, J., and Schroeder, A. (2018). Therapeutic nanoparticles penetrate leaves and deliver nutrients to agricultural crops. Scientific Reports 8, 1–10.

Khan, A., Numan, M., Khan, A.L., Lee, I.-J., Imran, M., Asaf, S., and Al-Harrasi, A. (2020). Melatonin: Awakening the defense mechanisms during plant oxidative stress. Plants 9, 407.

Khan, M.I.R., Fatma, M., Per, T.S., Anjum, N.A., and Khan, N.A. (2015). Salicylic acid-induced abiotic stress tolerance and underlying mechanisms in plants. Frontiers in plant science 6, 462.

Klokić, I., Koleška, I., Hasanagić, D., Murtić, S., Bosančić, B., and Todorović, V. (2020). Biostimulants’ influence on tomato fruit characteristics at conventional and low-input NPK regime. Acta Agriculturae Scandinavica, Section B—Soil & Plant Science 70, 233–240.

Knopf, R.R., Feder, A., Mayer, K., Lin, A., Rozenberg, M., Schaller, A., and Adam, Z. (2012). Rhomboid proteins in the chloroplast envelope affect the level of allene oxide synthase in Arabidopsis thaliana. The Plant Journal 72, 559–571.

Koressaar, T., and Remm, M. (2007). Enhancements and modifications of primer design program Primer3. Bioinformatics 23, 1289–1291.

Langmead, B., and Salzberg, S.L. (2012). Fast gapped-read alignment with Bowtie 2. Nature methods 9, 357.

Latif, S., Gurusinghe, S., Weston, P.A., Quinn, J.C., Piltz, J.W., and Weston, L.A. (2020). Metabolomic approaches for the identification of flavonoids associated with weed suppression in selected Hardseeded annual pasture legumes. Plant and Soil 447, 199–218.

Liu, S., Li, M., Su, L., Ge, K., Li, L., Li, X., Liu, X., and Li, L. (2016). Negative feedback regulation of ABA biosynthesis in peanut (Arachis hypogaea): a transcription factor complex inhibits AhNCED1 expression during water stress. Scientific reports 6, 1–11.

Lu, Y., and Yao, J. (2018). Chloroplasts at the crossroad of photosynthesis, pathogen infection and plant defense. International journal of molecular sciences 19, 3900.

Maksymiec, W., Wianowska, D., Dawidowicz, A.L., Radkiewicz, S., Mardarowicz, M., and Krupa, Z. (2005). The level of jasmonic acid in Arabidopsis thaliana and Phaseolus coccineus plants under heavy metal stress. Journal of plant physiology 162, 1338–1346.

Mátyás, K.K., Hegedűs, G., Taller, J., Farkas, E., Decsi, K., Kutasy, B., Kálmán, N., Nagy, E., Kolics, B., and Virág, E. (2019). Different expression pattern of flowering pathway genes contribute to male or female organ development during floral transition in the monoecious weed Ambrosia artemisiifolia L.(Asteraceae). PeerJ 7, e7421.

Mayer, L.D., Bally, M.B., Hope, M.J., and Cullis, P.R. (1986). Techniques for encapsulating bioactive agents into liposomes. Chemistry and physics of lipids 40, 333–345.

Moses, T., Papadopoulou, K.K., and Osbourn, A. (2014). Metabolic and functional diversity of saponins, biosynthetic intermediates and semi-synthetic derivatives. Critical reviews in biochemistry and molecular biology 49, 439–462.

Mou, Z., Fan, W., and Dong, X. (2003). Inducers of plant systemic acquired resistance regulate NPR1 function through redox changes. Cell 113, 935–944.

Mujeeb, F., Bajpai, P., and Pathak, N. (2014). Phytochemical evaluation, antimicrobial activity, and determination of bioactive components from leaves of Aegle marmelos. BioMed research international 2014.

Narwal, S. (2006). Allelopathy in ecological sustainable agriculture. In Allelopathy (Springer), pp. 537–564.

Narwal, S.S. (2012). Allelopathy in crop production. (Scientific publishers).

Noutoshi, Y., Okazaki, M., Kida, T., Nishina, Y., Morishita, Y., Ogawa, T., Suzuki, H., Shibata, D., Jikumaru, Y., and Hanada, A. (2012). Novel plant immune-priming compounds identified via high-throughput chemical screening target salicylic acid glucosyltransferases in Arabidopsis. The Plant Cell 24, 3795–3804.

Olusanmi, M., and Amadi, J. (2010). Studies on the antimicrobial properties and phytochemical screening of garlic (Allium sativum) extracts. Ethnobotanical leaflets 2009, 10.

Parađiković, N., Teklić, T., Zeljković, S., Lisjak, M., and Špoljarević, M. (2019). Biostimulants research in some horticultural plant species—A review. Food and Energy Security 8, e00162.

Pastor, V., Luna, E., Ton, J., Cerezo, M., García-Agustín, P., and Flors, V. (2013). Fine tuning of reactive oxygen species homeostasis regulates primed immune responses in Arabidopsis. Molecular plant-microbe interactions 26, 1334–1344.

Pieczul, K., Dobrzycka, A., Wolko, J., Perek, A., Zielezińska, M., Bocianowski, J., and Rybus-Zajac, M. (2020). The activity of β-glucosidase and guaiacol peroxidase in different genotypes of winter oilseed rape (Brassica napus L.) infected by Alternaria black spot fungi. Acta Physiologiae Plantarum 42, 1–9.

Prabha, D., Negi, S., Kumari, P., Negi, Y.K., and Chauhan, J. (2016). Effect of seed priming with some plant leaf extract on seedling growth characteristics and root rot disease in tomato. International Journal of Agriculture System 4, 46–51.

Reddy, A.R., Chaitanya, K., Jutur, P., and Sumithra, K. (2004). Differential antioxidative responses to water stress among five mulberry (Morus alba L.) cultivars. Environmental and experimental botany 52, 33–42.

Robinson, M.D., McCarthy, D.J., and Smyth, G.K. (2010). edgeR: a Bioconductor package for differential expression analysis of digital gene expression data. Bioinformatics 26, 139–140.

Ruan, J., Zhou, Y., Zhou, M., Yan, J., Khurshid, M., Weng, W., Cheng, J., and Zhang, K. (2019). Jasmonic acid signaling pathway in plants. International journal of molecular sciences 20, 2479.

Saez, G., Bannister, W., and Bannister, J. (1990). Free radicals and thiol compounds. The role of glutathione against free radical toxicity. (Boca Raton FL: CRC Press Inc).

Satish, S., Mohana, D., Ranhavendra, M., and Raveesha, K. (2007). Antifungal activity of some plant extracts against important seed borne pathogens of Aspergillus sp. Journal of Agricultural technology 3, 109–119.

Schwessinger, B., and Ronald, P.C. (2012). Plant innate immunity: perception of conserved microbial signatures. Annual review of plant biology 63, 451–482.

Sharma, H.S., Fleming, C., Selby, C., Rao, J., and Martin, T. (2014). Plant biostimulants: a review on the processing of macroalgae and use of extracts for crop management to reduce abiotic and biotic stresses. Journal of applied phycology 26, 465–490.

Silverman, P., Seskar, M., Kanter, D., Schweizer, P., Metraux, J.-P., and Raskin, I. (1995). Salicylic acid in rice (biosynthesis, conjugation, and possible role). Plant physiology 108, 633–639.

Singh, S., Kumar, V., Kapoor, D., Kumar, S., Singh, S., Dhanjal, D.S., Datta, S., Samuel, J., Dey, P., and Wang, S. (2020). Revealing on hydrogen sulfide and nitric oxide signals co-ordination for plant growth under stress conditions. Physiologia Plantarum 168, 301–317.

Sousa, R.H., Carvalho, F.E., Ribeiro, C.W., Passaia, G., Cunha, J.R., LIMA-MELO, Y., MARGIS-PINHEIRO, M., and Silveira, J.A. (2015). Peroxisomal APX knockdown triggers antioxidant mechanisms favourable for coping with high photorespiratory H 2 O 2 induced by CAT deficiency in rice. Plant, cell & environment 38, 499–513.

Špoljarević, M., Štolfa, I., Lisjak, M., Stanisavljević, A., Vinković, T., Agić, D., Parađiković, N., Teklić, T., Engler, M., and Klešić, K. (2010). Strawberry (Fragaria x ananassa Duch) leaf antioxidative response to biostimulators and reduced fertilization with N and K. Poljoprivreda 16, 50–56.

Taheri, P., and Tarighi, S. (2010). Riboflavin induces resistance in rice against Rhizoctonia solani via jasmonate-mediated priming of phenylpropanoid pathway. Journal of plant physiology 167, 201–208.

Thines, B., Katsir, L., Melotto, M., Niu, Y., Mandaokar, A., Liu, G., Nomura, K., He, S.Y., Howe, G.A., and Browse, J. (2007). JAZ repressor proteins are targets of the SCF COI1 complex during jasmonate signalling. Nature 448, 661–665.

Toscano, S., Romano, D., Massa, D., Bulgari, R., Franzoni, G., and Ferrante, A. (2018). Biostimulant applications in low input horticultural cultivation systems= I biostimolanti nei sistemi colturali ortofloricoli a basso impatto ambientale.

Ugena, L., Hýlová, A., Podlešáková, K., Humplík, J.F., Doležal, K., Diego, N.D., and Spíchal, L. (2018). Characterization of biostimulant mode of action using novel multi-trait high-throughput screening of Arabidopsis germination and rosette growth. Frontiers in plant science 9, 1327.

Untergasser, A., Cutcutache, I., Koressaar, T., Ye, J., Faircloth, B.C., Remm, M., and Rozen, S.G. (2012). Primer3—new capabilities and interfaces. Nucleic acids research 40, e115–e115.

Vanholme, B., Grunewald, W., Bateman, A., Kohchi, T., and Gheysen, G. (2007). The tify family previously known as ZIM. Trends in plant science 12, 239–244.

Vargas-Hernandez, M., Macias-Bobadilla, I., Guevara-Gonzalez, R.G., Romero-Gomez, S.d.J., Rico-Garcia, E., Ocampo-Velazquez, R.V., Alvarez-Arquieta, L.d.L., and Torres-Pacheco, I. (2017). Plant hormesis management with biostimulants of biotic origin in agriculture. Frontiers in plant science 8, 1762.

Virág, E., Hegedűs, G., Barta, E., Nagy, E., Mátyás, K., Kolics, B., and Taller, J. (2016). Illumina sequencing of common (short) ragweed (Ambrosia artemisiifolia L.) reproductive organs and leaves. Frontiers in plant science 7, 1506.

Walters, D.R., Newton, A.C., and Lyon, G.D. (2014). Induced resistance for plant defense: a sustainable approach to crop protection. (John Wiley & Sons).

Wang, W., Zhang, G., Yang, S., Zhang, J., Deng, Y., Qi, J., Wu, J., Fu, D., Wang, W., and Hao, Q. (2021). Overexpression of isochorismate synthase enhances drought tolerance in barley. Journal of Plant Physiology, 153404.

Wendehenne, D., Gao, Q.-m., Kachroo, A., and Kachroo, P. (2014). Free radical-mediated systemic immunity in plants. Current opinion in plant biology 20, 127–134.

Wildermuth, M.C., Dewdney, J., Wu, G., and Ausubel, F.M. (2001). Isochorismate synthase is required to synthesize salicylic acid for plant defence. Nature 414, 562–565.

Wu, G., Shortt, B.J., Lawrence, E.B., Leon, J., Fitzsimmons, K.C., Levine, E.B., Raskin, I., and Shah, D.M. (1997). Activation of host defense mechanisms by elevated production of H2O2 in transgenic plants. Plant Physiology 115, 427–435.

Zhang, Y., Fan, W., Kinkema, M., Li, X., and Dong, X. (1999). Interaction of NPR1 with basic leucine zipper protein transcription factors that bind sequences required for salicylic acid induction of the PR-1 gene. Proceedings of the National Academy of Sciences 96, 6523–6528.

